# Sedimentary factories and ecosystem change across the Permian-Triassic Critical Interval (P-TrCI) – insights from the Xiakou area (South China)

**DOI:** 10.1101/2020.08.10.244210

**Authors:** Yu Pei, Jan-Peter Duda, Joachim Reitner

## Abstract

The Permian-Triassic mass extinction included a potentially catastrophic decline of biodiversity, but ecosystem change across this event remains poorly characterized. Here we reconstruct sedimentary factories and ecosystem change across the Permian-Triassic Critical Interval (P-TrCI) in the Xiakou area (South China). Six microfacies (MF) were classified. The succession begins with a eukaryote-controlled carbonate factory (MF-1) that passes upward into an organomineralization-dominated carbonate factory (MF-2–3). Organic-rich marls atop these units reflect carbonate factory collapse (MF-4). Organomineralization-driven carbonate formation restarts prior to the Permian-Triassic boundary (MF-5) and subsequently develops into a mixed carbonate factory where organomineralization and biomineralization are almost equally important (MF-6). MF-1 reflects oxygenated shallow water environments. In contrast, MF-2–6 were all deposited in somewhat deeper environments, some of which episodically exhibited elevated salinities, oxygen depletion, and, possibly, euxinic conditions. Our results demonstrate that distinct changes in carbonate production styles, biodiversity, and environmental conditions are not synchronous at Xiakou. Furthermore, the Xiakou record is strikingly different to that of other localities, even from the same area (e.g., the Global Stratotype Section and Point section at Meishan). Together, these findings highlight the enormous complexity of the P-TrCI and calls simplified views of the Permian-Triassic mass extinction into question.

## 1 Introduction

The Permian-Triassic (P-Tr) mass extinction is characterized by a potentially catastrophic decline of biodiversity in marine and terrestrial ecosystems (Benton and Twitchett 2003; Wignall 2007; Chen and Benton 2012; Payne and Clapham 2012). For the marine realm, this view is mainly based on rarefaction curves that indicate as many as 96% of all species disappeared (Raup 1979) and findings from fossil databases that suggest 52% of all families went extinct (Sepkoski et al. 1981; Raup and Sepkoski 1982). Later studies based on paleontological databases supported this view (Fan et al. 2020), proposing that 78% of all marine genera were wiped out at the end of the Permian (Alroy et al. 2008). Others argue that these extinction rates might be overestimated (cf. Erwin 1994; Stanley 2016).

A prolonged period of recovery followed the P-Tr mass extinction (Lehrmann et al. 2006; Chen and Benton 2012). Strong shifts in the carbon stable isotopic composition of sedimentary carbonates (δ^13^C_carb_ of ca. -3–8‰) support the notion of long-lasting ecological disturbances in its aftermath (Payne et al. 2004). Marine sulfate concentrations were significantly lower in the Early Triassic as compared to today (≤ 6 mM, i.e., < 20% of modern-day seawater), as indicated by profound variations in carbonate associated sulfate and sulfate sulfur isotopes (δ^34^S_cas_ of ca. 9‰–44‰ and δ^34^S_sulfate_ of ca. 10‰–32‰, respectively) (Luo et al. 2010; Song et al. 2014; Bernasconi et al. 2017). Organisms in Early Triassic oceans were additionally stressed by high levels of toxic ammonium and a decline in bioavailable nitrogen, as indicated by analysis of bulk rock nitrogen isotopes (δ^15^N_bulk_ of ca. 1–3‰) (Grasby et al. 2019; Sun et al. 2019).

An important characteristic of the P-Tr mass extinction is the widespread occurrence of unusual sedimentary features in the event’s aftermath (Woods 2014; Chen et al. 2019). Examples include vermicular fabrics in limestones (Zhao et al. 2008) as well as fan- and calyx-shaped calcium carbonate precipitates (Woods et al. 1999; Woods 2014; Heindel et al. 2015). A further striking feature is the abundance of microbialites that developed on carbonate platforms in low-latitude regions of the Paleo-Tethys (Lehrmann 1999; Kershaw et al. 1999, 2012; Ezaki et al. 2003, 2008; Wang et al. 2005; Liu et al., 2007; Wu et al. 2007; Yang et al. 2011, 2019; Forel 2013; Lehrmann et al. 2015; Wang et al. 2016; Adachi et al. 2017; Fang et al. 2017; Tang et al. 2017; Wu et al. 2017; Chen et al. 2019; Foster et al. 2019; Pei et al. 2019; Wang et al. 2019). The unusual sedimentary features may reflect the persistence of unconventional environmental conditions associated with the end-Permian crisis (Bottjer et al. 2008; Chen and Benton 2012), while the widespread proliferation of microbial mats likely resulted from a suppressed ecological competition during this time (Foster et al. 2020).

The causes and triggers of the P-Tr mass extinction are still debated. One of Earths largest continental flood basalt provinces – the Siberian Traps – formed during this time and may have led to the mass extinction (Burgess and Bowring 2015; Burgess et al. 2017), as for instance indicated by a marked mercury anomaly (Wang et al. 2018). Volatiles such as halogens (Broadley et al. 2018), CO_2_, and SO_2_ (Wignall 2007) released from the massive, rapid outpouring of lavas could have affected Earth’s climate, raising seawater temperatures (Joachimski et al. 2012; Sun et al. 2012) and spreading anoxic conditions throughout Late Permian oceans (Grice et al. 2005; Brennecka et al. 2011; Elrick et al. 2017; Huang et al., 2017; Penn et al. 2018; Zhang et al. 2018). Other potential causes of the P-Tr mass extinction that were possibly, but not necessarily, related to volcanism include ocean acidification (Payne et al. 2010; Hinojosa et al. 2012; Clarkson et al. 2015) and hypercapnia (Knoll et al. 2007). It is likely that a combination of forces and processes interacted in complex ways to cause the mass extinction, and also make it difficult to directly link those causes to environmental consequences during this period.

This apocalyptic scenario might be oversimplified because fossil records covering the P-Tr extinction event are largely limited to marine shelf environments of Pangaea, reflecting conditions in a narrow belt and not necessarily those of remote parts of oceans. Furthermore, most of today’s organism families are unlikely to be preserved as fossils (Plotnick et al. 2016). By analogy, paleontological databases only comprise a fraction of groups that existed during P-Tr times. At the same time, fossilized groups are variably and inconsistently classified, making quantitative comparisons complicated. The occurrence of various Lazarus taxa (Jablonski 1986) in the Triassic (Erwin 1994; Wignall and Benton 1999) suggests the presence of unrecorded habitable refugia and of taphonomic biases, emphasizing the significance of such problems (Fraiser et al. 2011). A sound understanding of the complex interplay between biotic and abiotic processes through crucial junctures in Earth history requires more than studies of fossil databases (cf. Erwin 1994; Stanley 2016).

Rock-based approaches to studies of past ecosystems require well-preserved records. South China is known for numerous exquisitely preserved sedimentary successions that cover the Permian-Triassic critical transition, referred to as Permian-Triassic Critical Interval (P-TrCI) in the following. One well-known example is the Meishan Section – the Global Stratotype Section and Point (GSSP) for the P-Tr boundary (Yin et al. 2001). In this section, the P-Tr extinction horizon is positioned between Beds 24e-5 and 24e-6 (Chen et al. 2015), and it may be possible to directly correlate those beds with equivalent layers around the globe. A problem in making such correlations is that paleontological and δ^13^C_carb_ evidence from South China reveals a complex scenario of the P-TrCI because of the presence of one (Jin et al. 2000; Shen et al. 2018), two (Xie et al. 2007; Yin et al. 2012; Song et al. 2013; Chen et al. 2015), or even three extinction pulses (Yang et al. 1991). Inconsistencies between the manifestation of extinction events globally points out the need for detailed facies studies in order to better understand the variety of ecological changes during the P-TrCI.

Our study aims to unravel the complex interplay between biotic and abiotic processes through the P-TrCI – one of the most critical evolutionary junctures in Earth’s history – by studying facies in the Xiakou area (Hubei Province, South China) (Fig. 1). We use a rock-based approach that involves integration of sedimentary, paleontological, and biogeochemical evidence. Particular emphasis is placed on carbonate factory development during the P-TrCI as a measure of biological activity. This strategy allows for robust reconstruction of environmental conditions and ecosystem dynamics in this critical interval.

**Fig. 1.**
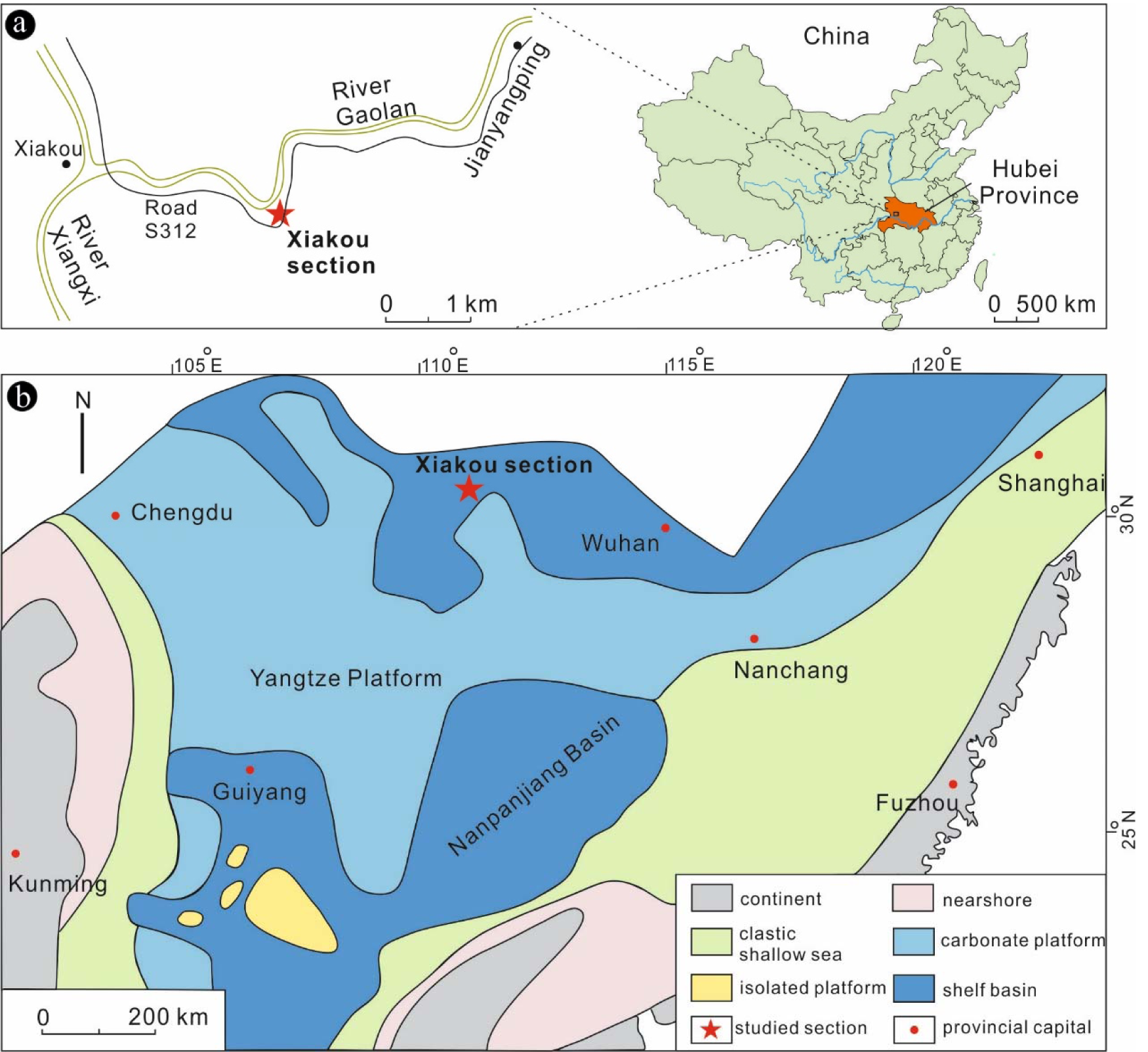
Study area. (a) Location of the Xiakou section, between the town of Xiakou and the village of Jianyangping (western Hubei Province, China). (b) Paleogeographical map of South China during the Changhsingian Stage (modified from Feng et al. 1997). The Xiakou area was situated in a shelf basin in the northernmost part of South China, adjacent to the Yangtze Platform

## 2 Geological background

The Xiakou section (GPS: N31°06.874′, E110°48.209′) is located between the town of Xiakou and the village of Jianyangping in western Hubei, South China (Fig. 1a). The section is well exposed and easily accessible in a gorge cut by the Gaolan River.

During P-Tr times, the Xiakou area was located in a shelf basin in the northernmost part of South China, adjacent to the Yangtze Platform (Feng et al. 1997) (Fig. 1b). The Xiakou section comprises (from base to top) the Changxing and Dalong Formations (both Changhsingian) as well as the Daye Formation (uppermost Changsinngian to Induan) (Fig. 2). The P-Tr boundary is placed within the lowermost Daye Formation at the base of Bed T_1_ (red line), as indicated by the first appearance of conodont *Hindeodus parvus* (Zhao et al. 2013).

**Fig. 2.**
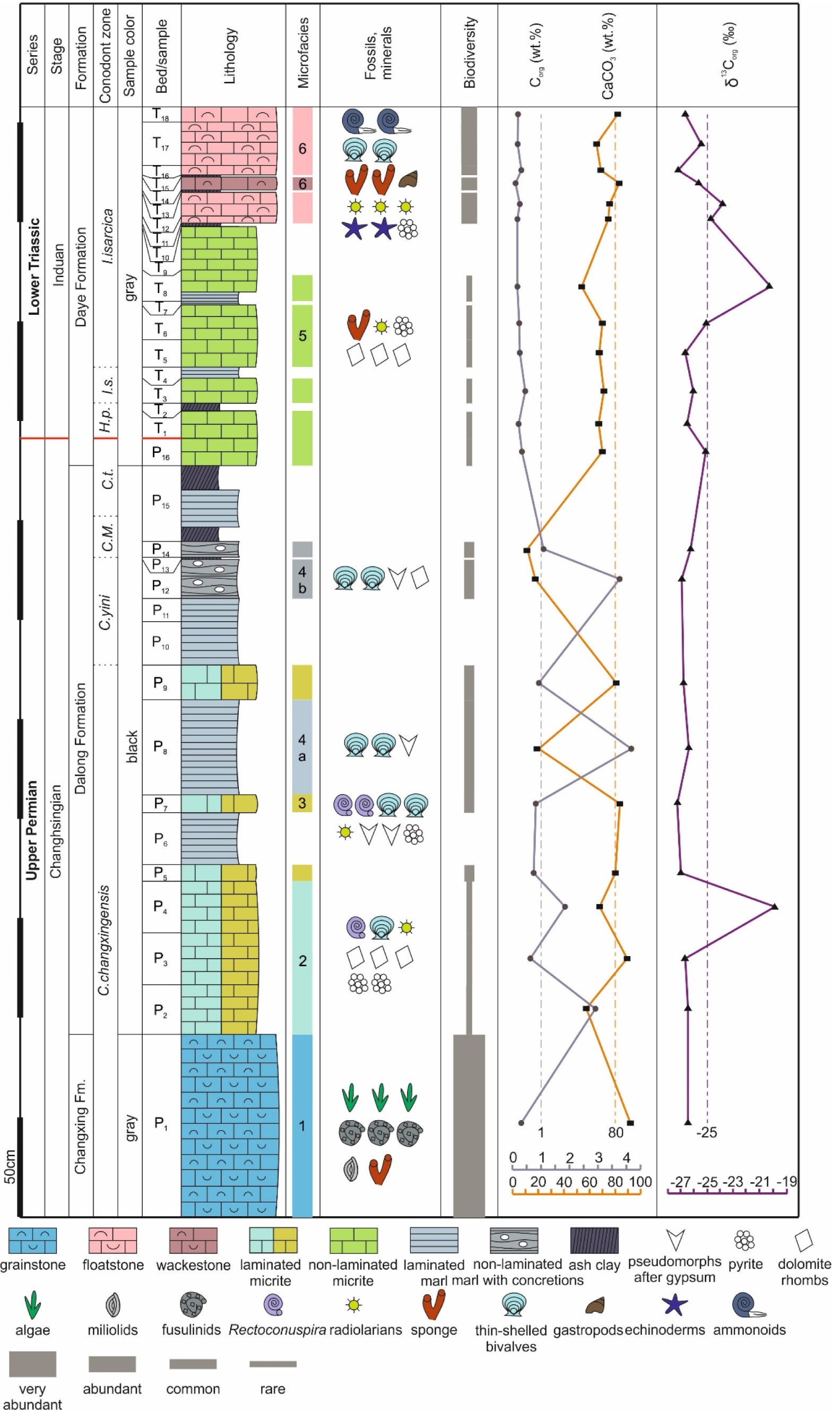
The Xiakou section, showing stratigraphical, sedimentological, paleontological, and geochemical features across the P-TrCI. Note that sample P_b1_ is not displayed. *C.M*. = *Clarkina Meishanensis, C.t. = Clarkina taylorae, H.p. = Hindeodus parvus, I.s. = Isarcicella staeschei*

## 3 Materials and methods

### 3.1 Fieldwork & petrography

Fieldwork was conducted in 2017 and 2019. Both campaigns included detailed observations and documentations of the Xiakou section and surrounding areas (Fig. 2). A total of 23 beds were sampled for subsequent petrographic and biogeochemical analyses (see below).

Thin sections (ca. three per sample) were analyzed using a Zeiss SteREO Discovery.V12 stereomicroscope and a Zeiss AXIO Imager.Z1 microscope coupled to an AxioCamMRc camera (transmitted and reflected light, respectively).

### 3.2 Analytical imaging techniques

For epifluorescence microscopy, a Zeiss AXIO Imager.Z1 microscope equipped with a high-pressure mercury arc lamp (HBO 50, Zeiss; controlled by an EBX 75 ISOLATED electronic transformer) and a 10 AF488 filter (excitation wavelength = BP 450–490 nm, emission wavelength = BP 515–565 nm) was used.

For cathodoluminescence (CL) microscopy, a Cambridge Instruments Citl CCL 8200 Mk3A cold-cathode system was linked to a Zeiss Axiolab microscope (operating voltage of ca. 15 kV; electric current of ca. 250–300 μA) and a Zeiss AxioCam 703 camera.

For field emission scanning electron microscopy (Fe-SEM), a Carl Zeiss LEO 1530 Gemini system was used. EDX spectra and elemental maps were acquired with an Oxford Instruments INCA x-act energy dispersive X-ray spectrometry (EDX) detector coupled to the Fe-SEM system.

Raman spectra were collected using a WITec alpha300R fiber-coupled ultra-high throughput spectrometer at the Geosciences Center, University of Göttingen. Before analysis, the system was calibrated using an integrated light source. The experimental setup included a 405 nm laser, 10 mW laser power, a 100x long working distance objective with a numerical aperture of 0.75, and a 1200 g mm^−1^ grating. This setup had a spectral resolution of 2.6 cm^−1^. The spectrometer was centered at 1530 cm^−1^, covering a spectral range from 122 cm^−1^ to 2759 cm^−1^. The 405 nm laser was chosen to reduce fluorescence effects. Each spectrum was collected by two accumulations, with an acquisition time of 60 s. Raman spectra were processed with the WITec project software. The background was subtracted using a rounded shape and band positions were determined by fitting a Lorentz function.

### 3.3 Bulk analyses

Bulk analyses (total organic carbon, C_org_; total inorganic carbon, C_carb_) were executed using a Leco RC612 carbon analyzer and a Hekatech Euro EA elemental analyzer. C_carb_ values were used to calculate CaCO_3_ contents for all samples.

### 3.4 Stable isotopes (δ^13^Ccarb, δ^18^Ocarb, δ^13^Corg)

Carbon and oxygen stable isotopic compositions of carbonates were analyzed in the Isotope Geology Department at the Geoscience Center of the Georg-August-Universität Göttingen (Germany). Individual mineral phases were sampled with a high-precision drill, permitting the isolation of minute amounts of sample powder (ca. 100–600 µg). The isotope measurements were performed at 70°C using a Thermo Scientific Kiel IV carbonate device coupled to a Finnigan DeltaPlus gas isotope mass spectrometer. Carbon and oxygen stable isotope ratios of carbonate minerals are reported as delta values relative to Vienna Pee Dee Belemnite (VPDB) reference standard (δ^13^C_carb_ and δ^18^O_carb_, respectively) Reproducibility was tested through the replicate analysis of standard NBS19 and was generally better than 0.1‰.

Carbon stable isotopic compositions of bulk organic matter were analyzed at the Centre for Stable Isotope Research and Analysis (KOSI) at the Georg-August-Universität Göttingen (Germany). An elemental analyzer (NA-2500 CE-Instruments) coupled to an isotope ratio mass spectrometer (Finnigan MAT Delta plus) was used to determine carbon isotopic abundances. About 1 g of each sample was decalcified, washed, and neutralized, and ca. 0.2–15 mg of the remaining material were analyzed. The carbon stable isotope ratios of bulk organic matter are reported as delta values relative to VPDB reference standard (δ^13^C_org_). For internal calibration an acetanilide standard was used (δ^13^C = –29.6 ‰; SD =0.1). The average δ^13^C_org_ value had a standard deviation of 0.3.

## 4 Results

### 4.1 Field observations

The part of the Xiakou section we investigated is about 9 meters thick (Fig. 2). It begins with ca. 4 meters of the uppermost Changxing Formation at the base (Beds P_b1_, and P_1_), which is composed of gray, thick-bedded to massive grainstones that locally contain silicate nodules. The overlying Dalong Formation (Beds P_2_–P_16_: Fig. 3a) is ca. 3 meters thick and mainly composed of black, thin-to medium-bedded micrites intercalated with black mudstones and greenish-yellowish volcanic ash layers (particularly abundant in Bed P_15_). The Daye Formation (only ca. 2 meters covered here, Beds T_1_–T_18_: Fig. 3a–b) terminates the section and consists of gray, thin-to medium-bedded micrites, wackestones, and floatstones, locally intercalated with black shales and thin greenish-yellowish volcanic ash layers. Few beds seem to pinch out laterally (T_5_, T_6_: Fig. 3b).

**Fig. 3.**
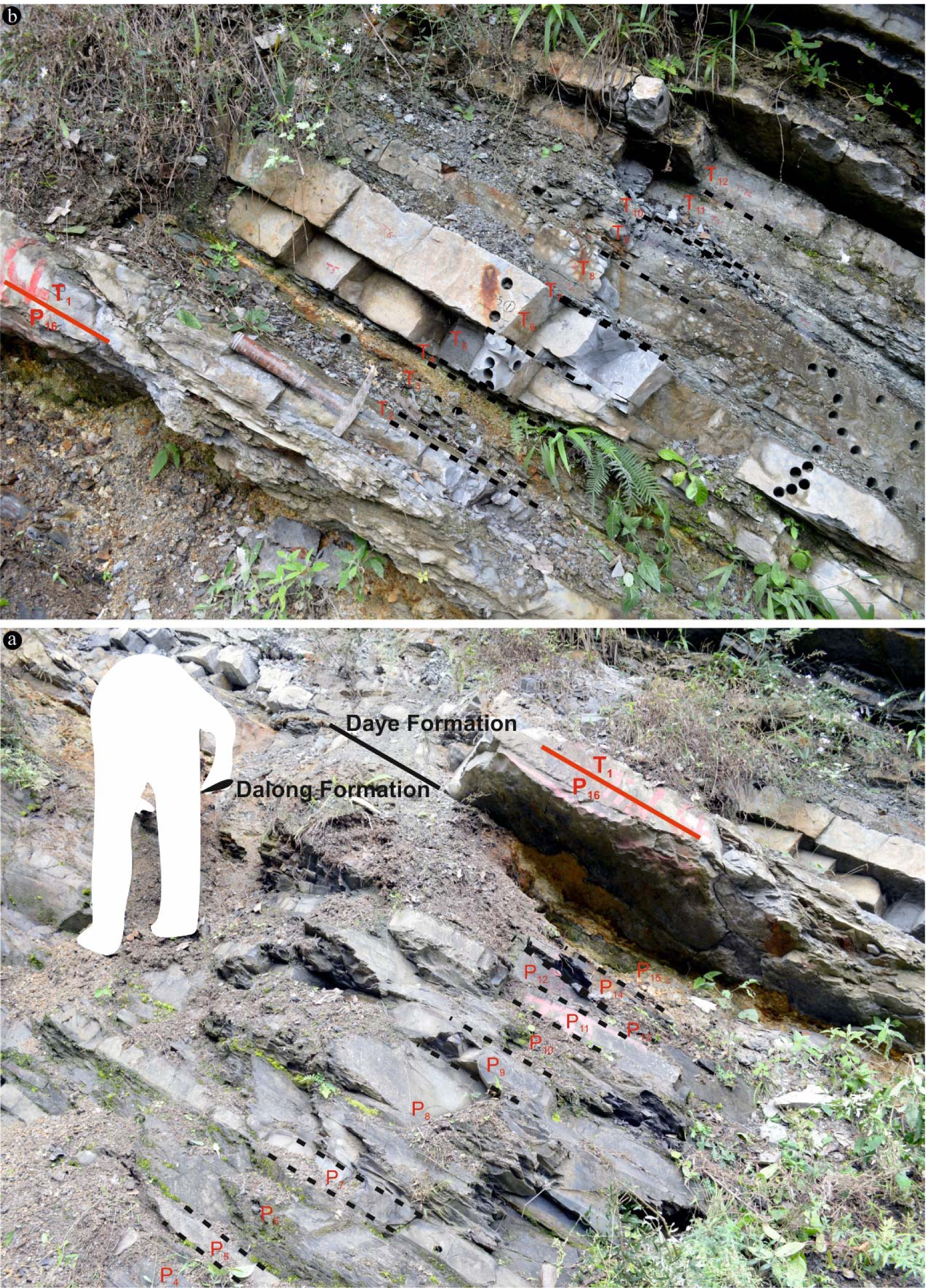
Field photos of the Xiakou section, showing the Dalong and Daye formations (beds P_4_–P_15_ and P_16_–T_1_, respectively). The P-Tr boundary is situated between beds P_16_ and T_1_ in the lowermost Daye Formation. Person (a) and hammer (b) for scale

### 4.2 Microfacies analysis

Based on sedimentary and paleontological criteria, six distinct microfacies (MF) can be distinguished.

#### 4.2.1 MF-1 Gray grainstone with dasyclad green algae and fusulinid foraminifers (Fig. 4a)

Samples: P_b1_, P_1_ (Changxing Formation)

**Fig. 4.**
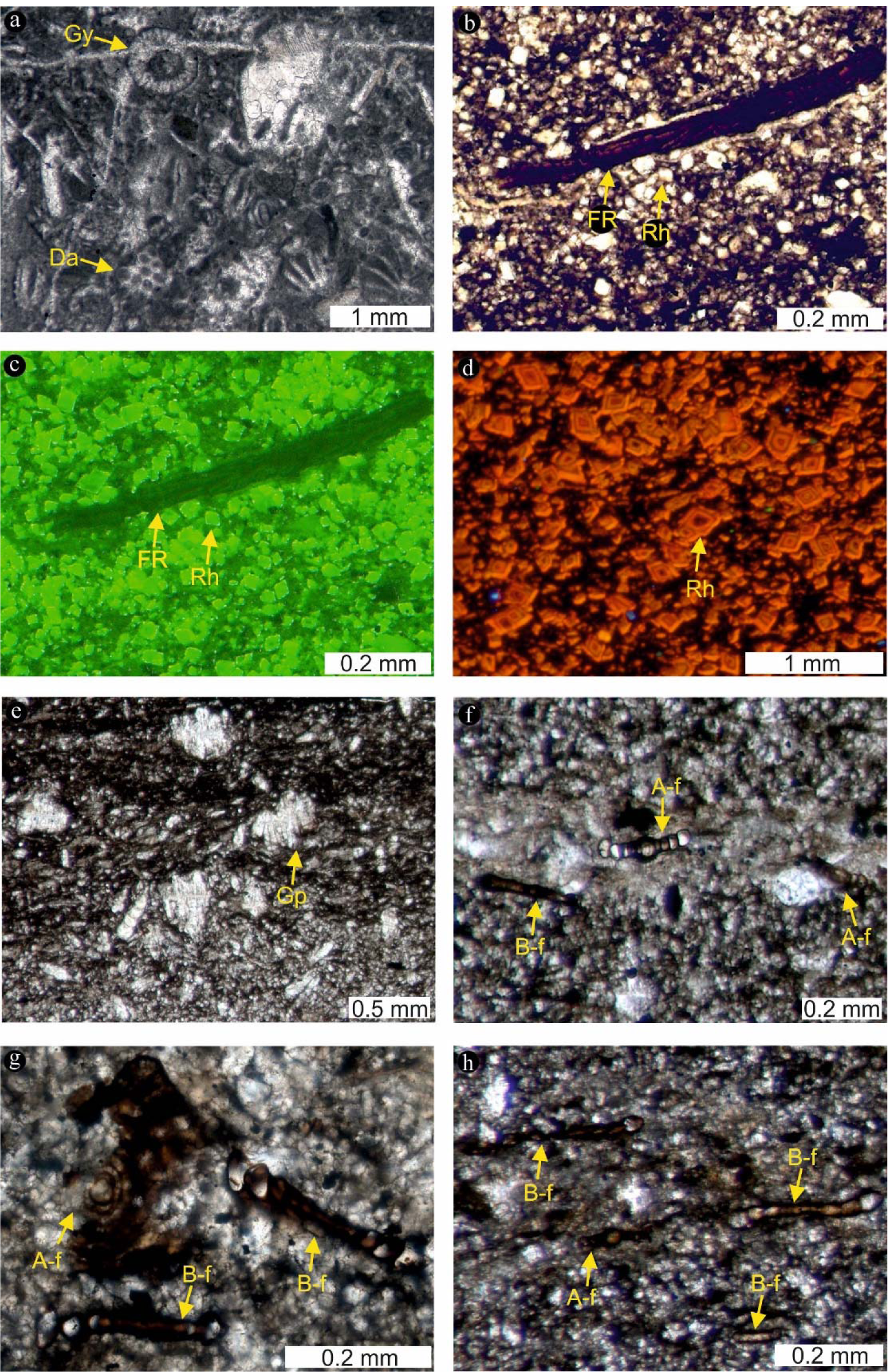
Thin section images of microfacies (MF) 1-3. (a) MF-1: Gray grainstone with dasyclad green algae (Da) and fusulinid foraminifers. Gy = Gymnocodiaceae. (b–d) MF-2: Black laminated micrite with dolomite rhombs (Rh). The dolomite rhombs (b) exhibit a strong green fluorescence (c; same thin section as b) and CL zoning (d). Fossils are rare in this MF and include fish remains (FR) (b–c). (e–h) MF-3: Black laminated micrite with abundant calcite pseudomorphs after gypsum and *Rectocornuspira* foraminifers. The calcite pseudomorphs after gypsum (Gp) locally form aggregates (e). Notably, this MF contains abundant A- and B-forms of *Rectocornuspira* (A-f and B-f in e–h, respectively). Both forms can clearly be distinguished based on the size of the initial chamber (large in A-form, small in B-form)

Carbonate components (mainly fossils) typically range in size from 0.5 mm to 2 mm. They are poorly rounded and moderately sorted. Few fossils are fragmented. The most abundant fossils are Dasycladaceae (e.g., *Mizzia*) and Fusulinidae. Other abundant constituents are Gymnocodiaceae and Miliolidae. In addition, demosponges, ostracods, echinoderms, bryozoans, brachiopods, and bivalves occur.

#### 4.2.2 MF-2 Black laminated micrite with dolomite rhombs (Fig. 4b–d)

Samples: P_2_–P_4_ (Dalong Formation)

The MF contains abundant small (5–25 µm) dolomite rhombs situated in a fine-grained calcite matrix. The rhombs exhibit a strong green fluorescence (especially in their outer rims) and CL zoning (Fig. 4b–d). In addition, there are rare occurrences of lenticular twinned calcite pseudomorphs (about 0.1 mm) after gypsum.

Fossils include thin-shelled bivalves, small benthic foraminifera, and fish bones (Fig. 4b). Locally, globular silica tests of radiolarians replaced by carbonate are observed.

#### 4.2.3 MF-3 Black laminated micrite with abundant calcite pseudomorphs after gypsum and Rectocornuspira foraminifers (Fig. 4e–h)

Samples: P_5_, P_7_, P_9_ (Dalong Formation)

This MF contains high abundances of calcite pseudomorphs after gypsum (0.1–0.5 mm; ca. 30% of total area in cross sections) (Fig. 4e). These components are lenticular and partly twinned and aggregated (Fig. 4e). In cross sections, about 64% of the pseudomorphs (N = 129) exhibit angles of 60°–120° between their long main axis and the sedimentary bedding plane (Fig. 5).

**Fig. 5.**
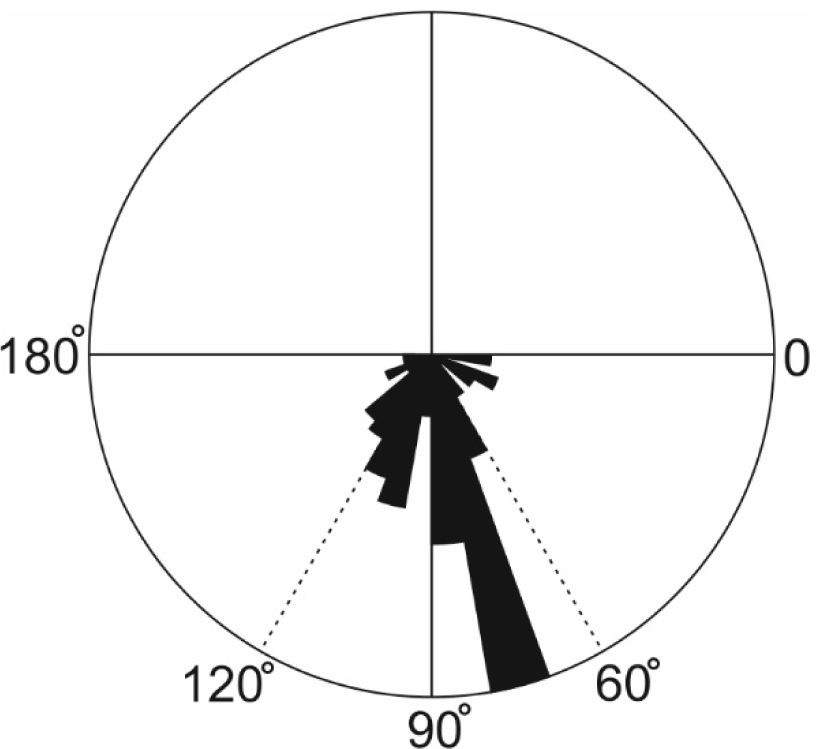
Rose diagram showing angles between long axes of lenticular-shaped pseudomorphs after gypsum and sedimentary bedding planes (measured in cross sections; N= 129). Note that about 64% of the angles range between 60° and 120°, suggesting that the gypsum crystals formed *in-situ*

Some calcite-rich laminae contain abundant megalospheric and microspheric stages of the benthic foraminifer *Rectocornuspira* (Fig. 4f–h), which are characterized by large and small initial chambers, respectively (Goldstein 1999). The megalospheric A-forms exhibit mean diameters of ca. 153 µm (N = 38), while the microspheric B-forms are ca. 249 µm in diameter on average (N = 75) (Fig. 6). Other fossils such as small benthic foraminifera, thin-shelled bivalves, and radiolarians were rarely observed.

**Fig. 6.**
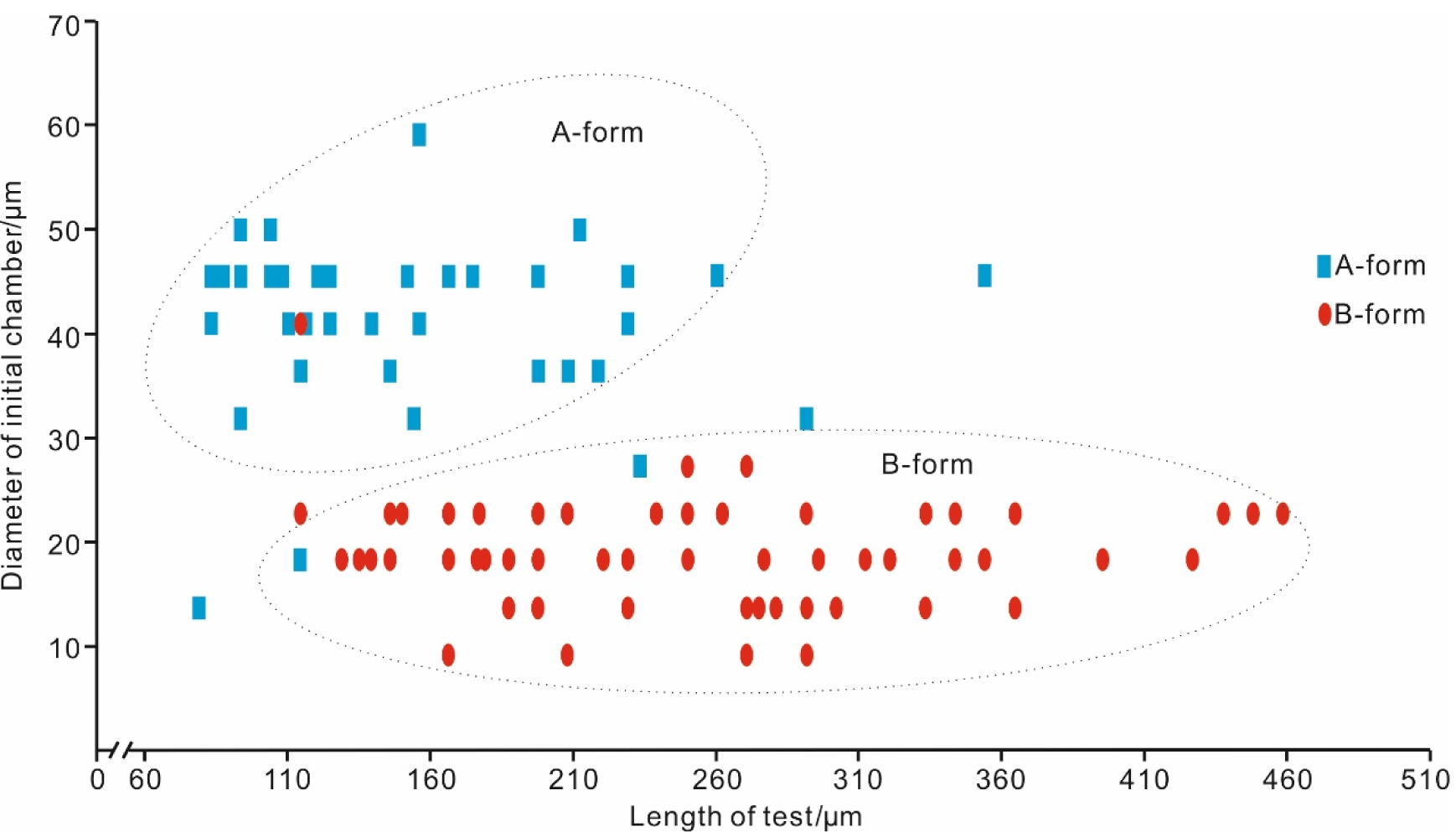
Crossplot diagram of morphological parameters of *Rectocornuspira* specimens in MF-3 (length of test vs diameter of initial chamber; N = 113). Note that specimens plot in two separate groups, supporting the interpretation as megalospheric and microspheric stages (A- and B-form, respectively)

#### 4.2.4 MF-4 Black marl

Samples: P_8_, P_12_, P_14_ (Dalong Formation)

This MF can be divided into two sub-types (MF-4a and MF-4-b; see below).

##### 4.2.4.1 MF-4a Black laminated marl with abundant fossil debris (Fig. 7a–b)

Sample: P_8_ (Dalong Formation)

**Fig. 7.**
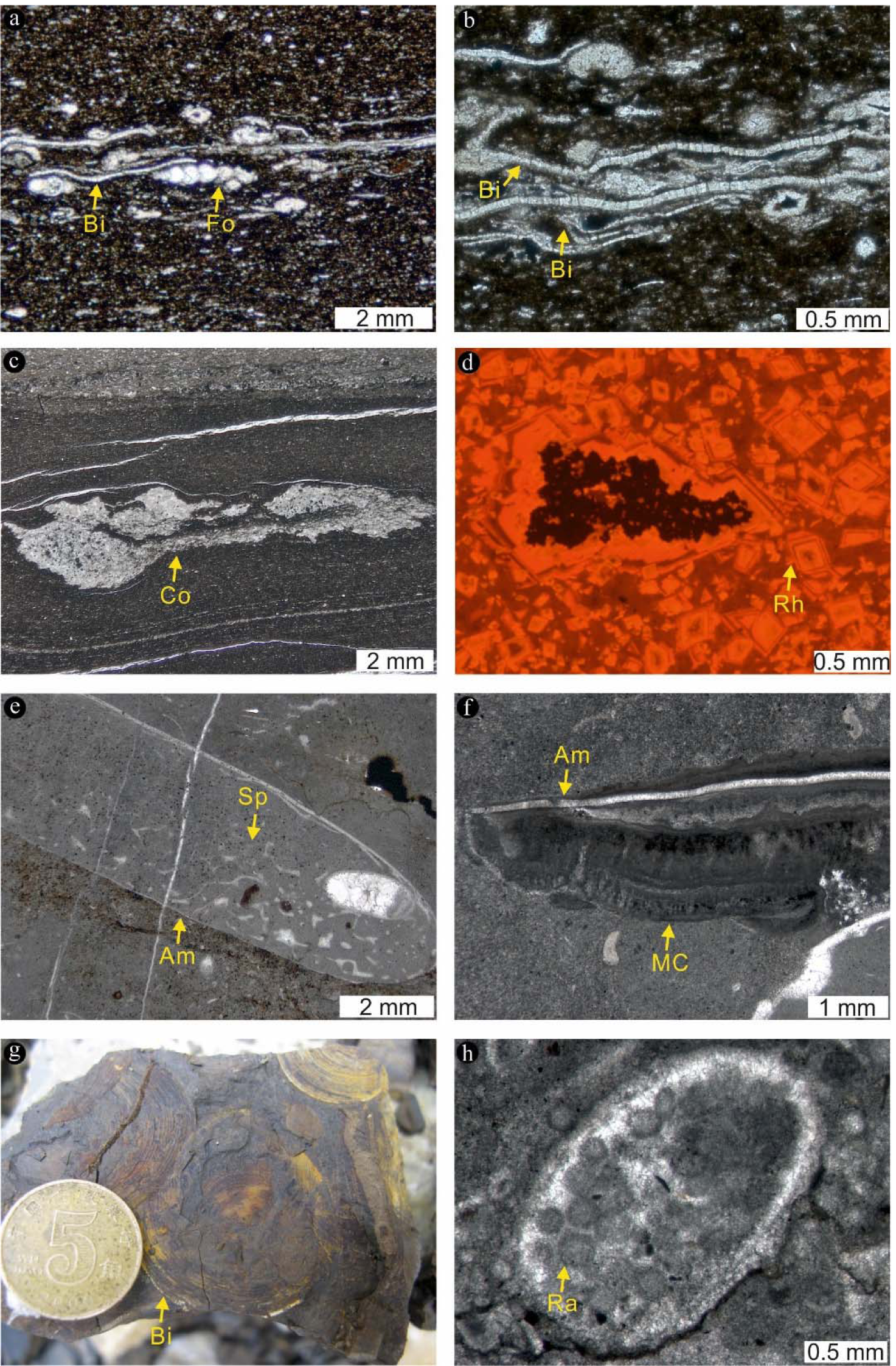
Thin section images of microfacies (MF) 4-6. (a–b) MF-4a: Black laminated marl with abundant fossil debris. Some layers contain well-preserved thin-shelled bivalves (Bi) and small benthic foraminifera (Fo) with hyaline radial tests (a). Other important features include intact bivalve fossils (Bi: b). (c) MF-4b: Black non-laminated marl with calcite and pyrite concretions (Co). (d) MF-5: Gray non-laminated micrite with dolomite rhombs (Rh). Rhombohedral-shaped crystals show similar CL characteristics as those in MF-2. (e–h) MF-6: Gray float-to wackestone with ammonoids (Am). Note that ammonoid shells served as substrates for non-spicular demosponges (Sp: e) and biofilms (as indicated by microbial calcite crusts, MC: f). Other abundant components include thin-shelled bivalves (Bi: g), gastropods, ostracods and radiolarians. Radiolarians (Ra) are locally densely packed (h)

**Fig. 8.**
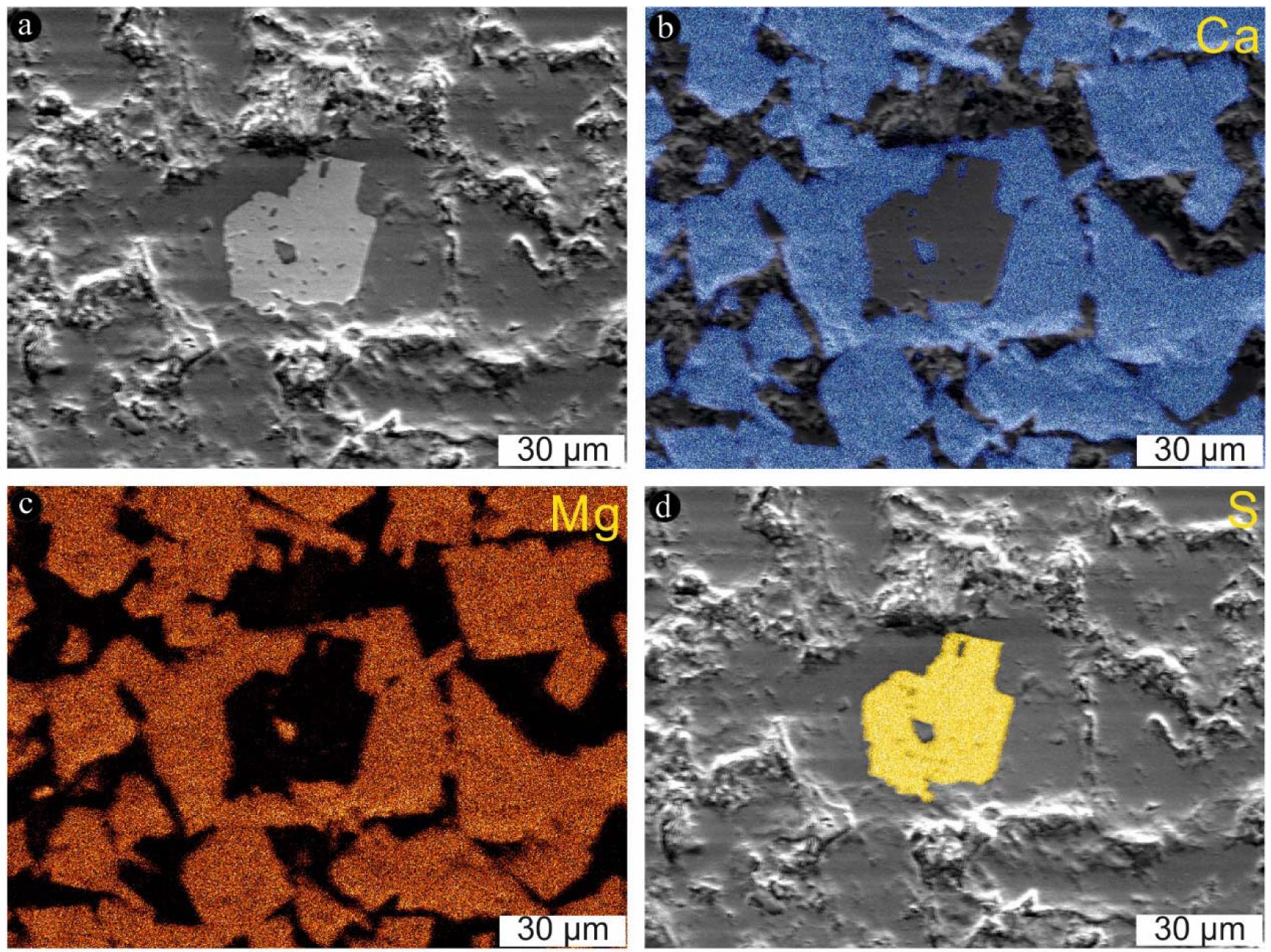
Field emission scanning electron microscopy – X-ray spectrometry (Fe-SEM EDX) images of dolomite rhombs in MF-5. (a) SEM image. (b) Calcium distribution. (c) Magnesium distribution. (d) Sulfur distribution

This MF subtype contains abundant fossil debris (0.01–0.1 mm) floating in a fine-grained matrix. Low C_carb_ and calculated CaCO_3_ contents (2.46 wt. % and 20.5 wt. %, respectively: Table 1) indicate that the matrix consists of carbonate and clay. Well-preserved thin-shelled bivalves and small benthic foraminifera with hyaline radial tests are enriched in certain layers (Fig. 7a). Some of the bivalve fossils in such layers are still intact (i.e., composed of both valves) (Fig. 7b). Few lenticular and partly twinned calcite pseudomorphs after gypsum are preserved. Furthermore, the MF contains small (25–100 µm) silicate mineral crystals.

**Table 1.**
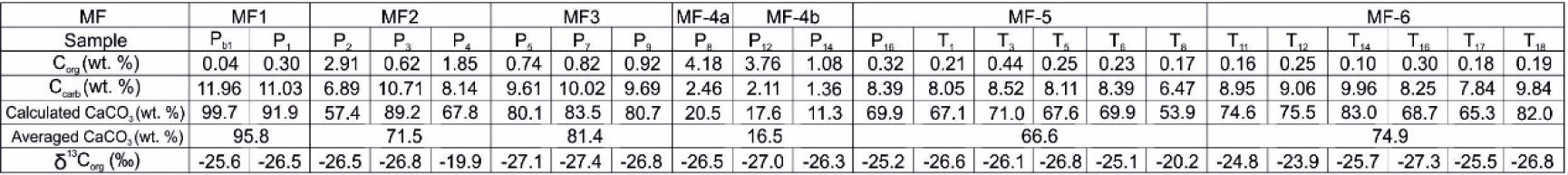
Bulk geochemical data, including carbon stable isotope signatures of organic matter (δ^13^C_org_)

##### 4.2.4.2 MF-4b Black non-laminated marl with calcite and pyrite concretions (Fig. 7c)

Samples: P_12_, P_14_ (Dalong Formation)

This MF subtype contains abundant 0.2–12 mm sized concretions composed of variable mixtures of calcite and pyrite in a fine-grained matrix (Fig. 7c). Low C_carb_ and calculated CaCO_3_ contents (≤2.11 wt. % and ≤17.6 wt. %, respectively: Table 1) indicate that the matrix consists of carbonate and clay. The concretions contain clasts, including dolomite rhombs. Abundant fragmented thin-shelled bivalves and small benthic foraminifera are observed. In addition, few fish remains and calcite pseudomorphs after gypsum occur. Furthermore, the MF contains small (25–100 µm) silicate mineral crystals.

#### 4.2.5 MF-5 Gray non-laminated micrite with dolomite rhombs (Fig. 7d)

Samples: P_16_ (Dalong Formation); T_1_, T_3_, T_5_, T_6_, T_8_ (Daye Formation)

This MF contains small (5–50µm) rhombohedral-shaped dolomite crystals (Figs. 7d, 8, 9) that show similar fluorescence and CL characteristics as those in MF-2 (Fig. 4d). Raman spectroscopy revealed that some of the dolomite rhombs (Fig. 9a–b) exhibit pyrite cores (Fig. 9a, c), which in turn encapsulate dolomite crystals (Fig. 9a, d). The MF contains a few specimens of *Earlandia*. A further feature are centimeter-sized concretions that encapsulate non-spicular demosponges and radiolarians. Notably, some of the concretions are surrounded by microbial calcite crusts.

**Fig. 9.**
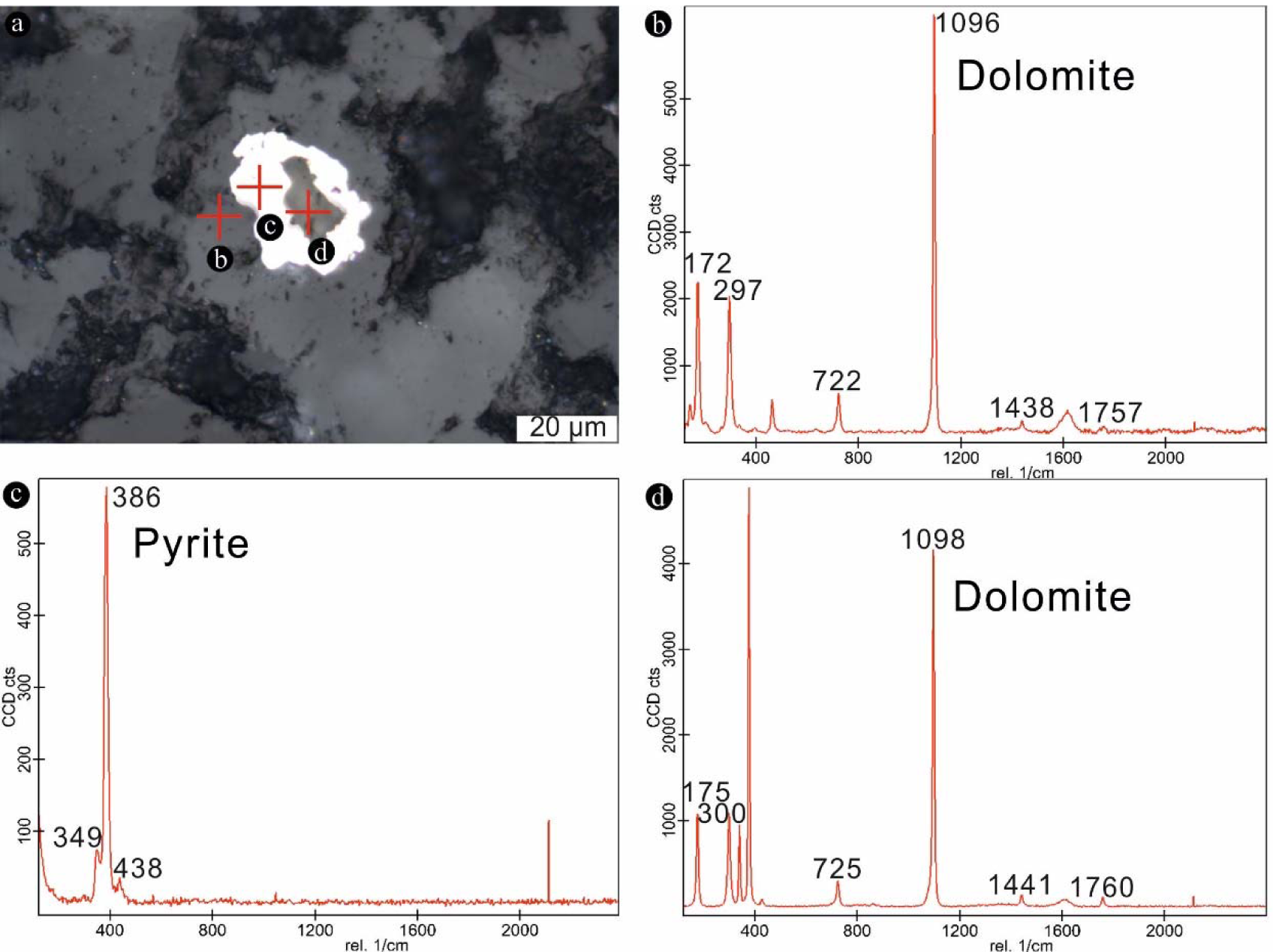
Raman spectroscopic features of dolomite rhombs in MF-5. (a) Positions of point measurements shown in b–d (reflected light image). Note that the dolomite rhombs (b) exhibit pyrite cores (c), that in turn encapsulate dolomite (d)

#### 4.2.6 MF-6 Gray float to wackestone with ammonoids (Fig. 7e–h)

Samples: T_11_, T_12_, T_14_, T_16_, T_17_, T_18_ (Daye Formation)

Carbonate components (mainly fossils) are typically >0.1 mm in diameter. They are poorly rounded and moderately sorted. The most abundant fossils are ammonoids (Fig. 7e). Shells of these organisms provided substrates for non-spicular demosponges (Fig. 7e) and microbial calcite crusts (Fig. 7f). Other abundant components are thin-shelled bivalves (Fig. 7g), gastropods, ostracods and radiolarians, the latter being locally densely packed (Fig. 7h).

### 4.3 Bulk geochemical characterization

The samples exhibit highly variable C_org_ contents that range between 0.04 wt.% and 4.18 wt.% (Fig. 2; Table 1). The highest C_org_ contents are observed in samples grouping into the MF-2 (P_2_ = 2.91 wt.%) and MF-4 (P_12_ = 3.76 wt.% and P_8_ = 4.18 wt.%). C_carb_ contents also vary profoundly, with values ranging from 1.36 wt.% to 11.96 wt.% (corresponding to calculated CaCO_3_ contents of 11.3 wt.% to 99.7 wt.%: Fig. 2; Table 1). C_org_ and CaCO_3_ contents are negatively correlated (Fig. 2; Table 1).

### 4.4 Stable isotopes (δ^13^Ccarb, δ^18^Ocarb, δ^13^Corg)

δ^13^C_carb_ values of different mineral phases vary from 0.3‰ to 4.2‰ (Fig. 10; Table 2). δ^18^O_carb_ values range from -9.2‰ to -1.6‰ (Fig. 10; Table 2). δ^13^C_carb_ and δ^18^O_carb_ do not exhibit a linear relationship, pointing against meteoric diagenesis (Bishop et al. 2014). A later fracture cement has a δ^13^C_carb_ value of 4.8‰ and a δ^18^O_carb_ value of -6.9‰. The MF show systematically different δ^13^C_carb_ and δ^18^O_carb_ values (Fig. 10; Table 2).

**Table 2.**
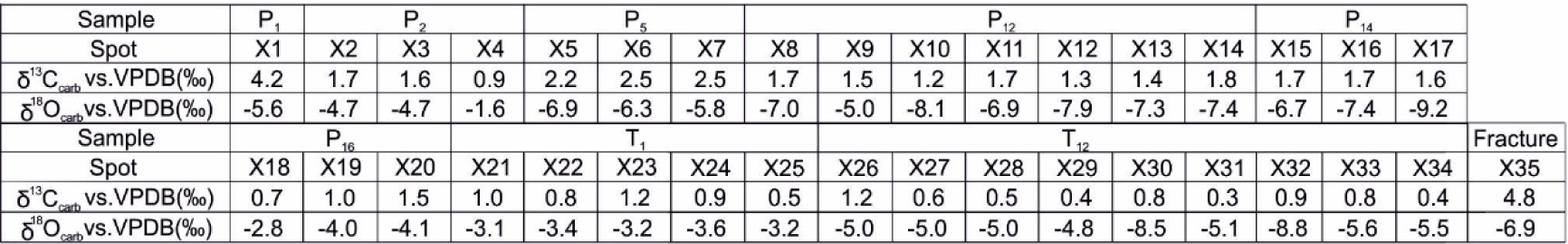
Carbon stable isotope and oxygen stable isotope signatures of carbonates (δ^13^C_carb,_ δ^18^O_carb_)

**Fig. 10.**
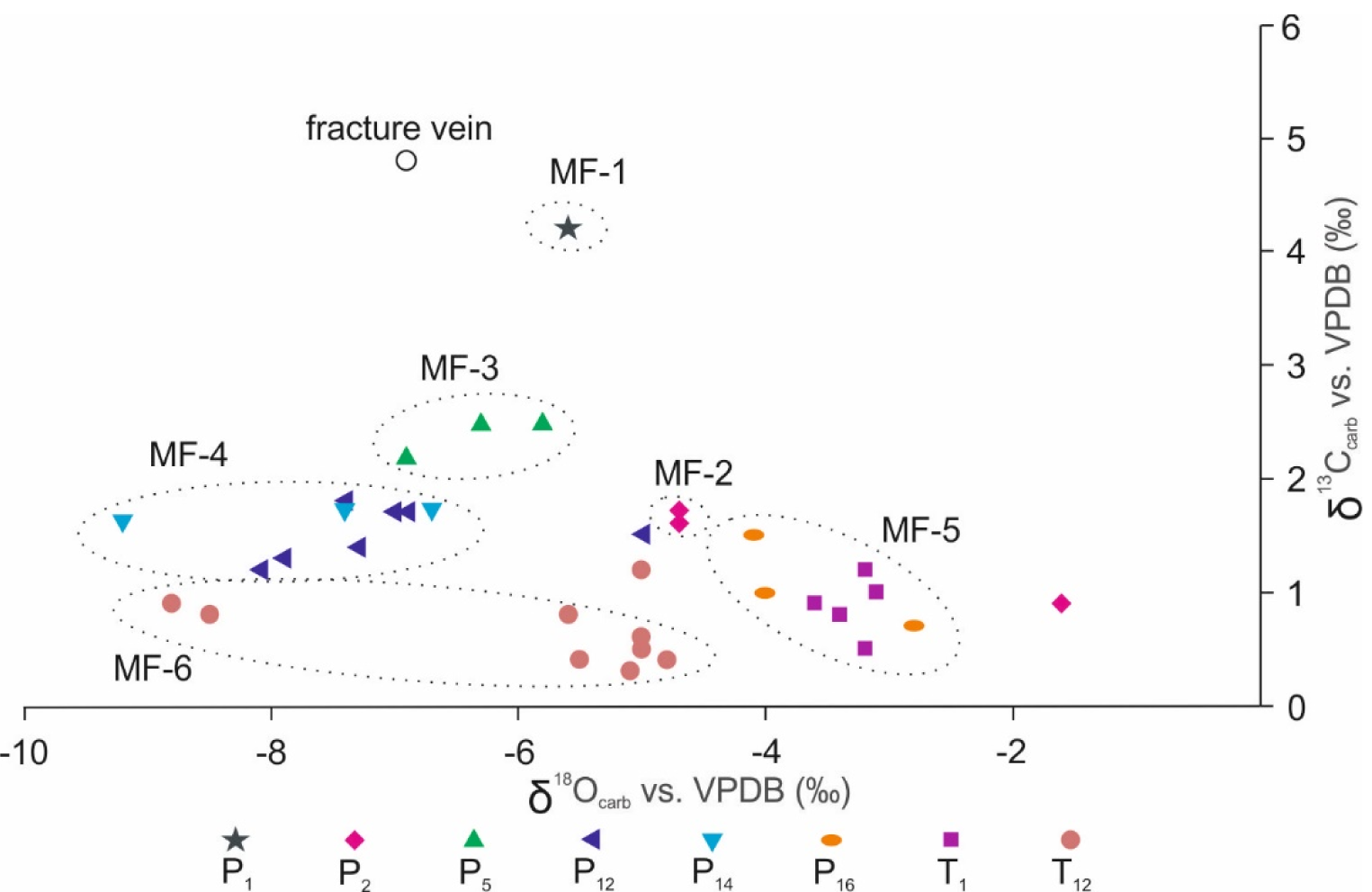
Carbon stable isotope and oxygen stable isotope data for individual carbonate phases in different samples. Note that the MF show systematically different δ^13^C_carb_ and δ^18^O_carb_ values (circled areas)

δ^13^C_org_ signatures of the samples vary between –27.4‰ and –19.9‰ (Fig. 2; Table 1). Most values cluster around ca. –25‰, but two samples show distinctly heavier signatures (P_4_ = –19.9‰, T_8_ = –20.2‰: Fig. 2; Table 1).

## 5 Discussion

### 5.1 Paleocommunities and -environments

Assemblages of calcareous algae (e.g., Dasycladaceae, Gymnocodiaceae) and photosymbiont-bearing foraminifera (e.g., Fusulinidae) indicate that MF-1 was deposited in oxygenated shallow water environments under tropical to subtropical conditions (Hottinger 1997; Langer and Hottinger 2000; Flügel 2010) (Figs. 4a, 9). In such environments, oxygenic phototrophs typically dominate primary productivity, which is consistent with δ^13^C_org_ values of about –26‰ (Fig. 2; Table 1). A significant uptake of ^12^C by autotrophic primary producers usually results in a concurrent enrichment of ^13^C in the water body, which is well in line with the high δ^13^C_carb_ value of 4.2‰ (Fig. 10; Table 2).

Occurrences of planktonic radiolarians and nektonic ammonoids in MF-2–3 and MF-5–6 (Figs. 7e, h, 11) suggest somewhat greater water depths than for MF-1. This observation is in good agreement with the presence of intact, thin-shelled bivalves that were preserved in gravity flow deposits (MF-4a) (Figs. 7b, 11) and indicate a relatively deep, low-energy environment. The presence of low-oxygen-tolerant and potentially chemosymbiont-bearing organisms such as non-spicular demosponges and thin-shelled bivalves such as *Claraia* in MF-2–6 (see Wignall and Hallam 1992; Hoffmann et al. 2005; McRoberts 2010; Zhao et al. 2013; Mills et al. 2014; Huang et al. 2018; Figs. 7a–b, e, g, 11) possibly reflect dysoxic to anoxic conditions at the seafloor. In the light of these findings, δ^13^C_org_ values of about –20‰ in beds P_4_ and T_8_ (Fig. 2; Table 1) might fingerprint significant contributions of organic matter by anoxygenic phototrophic bacteria to the bulk biomass (Preuß et al. 1989; Posth et al. 2017).

Porcelaneous Miliolina (e.g., abundant *Rectocornuspira* in MF-3, rare *Earlandia* in MF-5: Figs. 4f, 11) are ecologic opportunists commonly related to hypersaline conditions (Hallam and Wignall 1997; Groves and Altiner 2005; Flügel 2010). Elevated salinities are possibly also indicated by the presence of calcite pseudomorphs after gypsum (highly abundant in MF-3, rare in MF-2 and -4: Figs. 4e). About 64% of the calcite pseudomorphs exhibit angles of 60°–120° between their long main axis and the sedimentary bedding plane (N = 129; Fig. 5), indicating that gypsum formed *in situ* (Warren 2006). Gypsum has also been observed in P-TrCI strata of the Meishan section in South China (Liang 2002; Kaiho et al. 2006; Chen et al. 2015; Li and Jones 2017). Alternatively, the gypsum may be formed through the re-oxidation of hydrogen sulfide, as observed in organic matter or methane rich sediments affected by a pronounced sulfate reduction (Lin et al. 2016). Furthermore, potential biomarkers of halophilic Archaea were found in P-Tr boundary microbialites, suggesting that hypersaline conditions existed episodically in the upper water column of the Neo-Tethys (Heindel et al. 2018). Taken together, these findings indicate elevated salinities in parts of the Neo- and Paleo-Tethys, including in the Xiakou area.

### 5.2 Sedimentary factories

Carbonate production has been linked to biological processes throughout Earth’s history (James and Ginsburg 1979; Flügel 2010). Carbonate rocks can be formed from the accumulation of the skeletons of eukaryotic organisms. This process thus depends on the controlled precipitation of inorganic minerals by living organisms; that is, “biomineralization” *sensu stricto* (Mann 2002). Carbonate rocks can also form by the microbially induced precipitation of calcium carbonate. This process is related to chemical changes induced by metabolic activity, and commonly linked to exopolymeric substances (EPS) formed by microbial communities (Arp et al. 2001; Reitner et al. 2001; Riding 2010).

Carbonate precipitation can also be induced by organic matter (i.e., “organomineralization”). Typical products are autochthonous fine-grained carbonate (“automicrite”) and/or rhombohedral crystals (Addadi and Weiner 1985; Reitner 1993; Reitner et al. 1995a, b, 2000, 2001; Trichet and Défarge 1995). Automicrite is easily confused with micrite consisting of allochthonous carbonate material (“allomicrite”) but can commonly be distinguished by specific features (e.g., clotted and mottled fabrics, peloidal microstructures, strong fluorescence) (Wolf 1965; Reitner 1993; Reitner et al. 1995b). Tightly packed dolomite rhombs showing CL zoning are usually interpreted as diagenetic cements (Flügel 2010). However, “organomineralization” can also form rhombs (Addadi and Weiner 1985; Trichet and Défarge 1995). The observed organic matter enrichments in the rims of the dolomite rhombs (Fig. 4c) might imply that organic templates influenced crystal growth. In this view, the dolomite rhombs at Xiakou could be considered organominerals.

The investigated succession begins with a eukaryote-controlled carbonate factory, where the remains of shell-forming organisms significantly contribute to the sediment (MF-1: Figs. 4a, 11, 12a; Table 1). Automicrite is relatively rare and mainly restricted to taphonomic degradation of organisms such as sponges. In the section immediately above, the eukaryote-controlled sediment factory declined and was gradually replaced by fine-grained carbonate with abundant rhombohedral dolomite crystals (MF-2: Figs. 4b–d, 12b). The fine-grained matrix of MF-2 and MF-3 may also be automicrite. Moving upward in the section, the organomineralization-based carbonate factory collapses, as reflected in distinct black marls with only 16.5 wt.% carbonate (MF-4: Fig. 12b; Table 1). Following this interval, and just below the P-Tr boundary, carbonate formation started again. These carbonates (Bed P_16_) contain abundant rhombohedral dolomite crystals that are similar to ones found in MF-2 and also interpreted as organominerals (Figs. 7d, 12b; Table 1). This facies then passes into a mixed carbonate factory, where organomineralization and biomineralization are almost equally important (Figs. 7e–h, 11, 12c; Table 1).

**Fig. 11.**
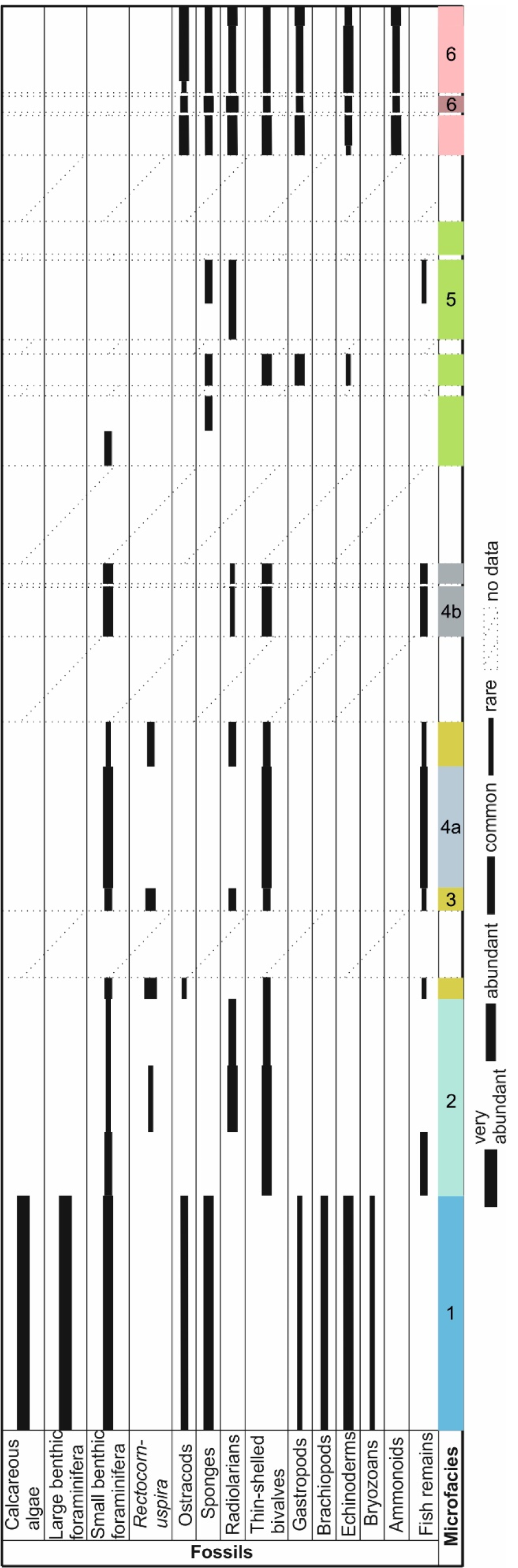
Distribution of fossils in different microfacies from the Xiakou section

**Fig. 12.**
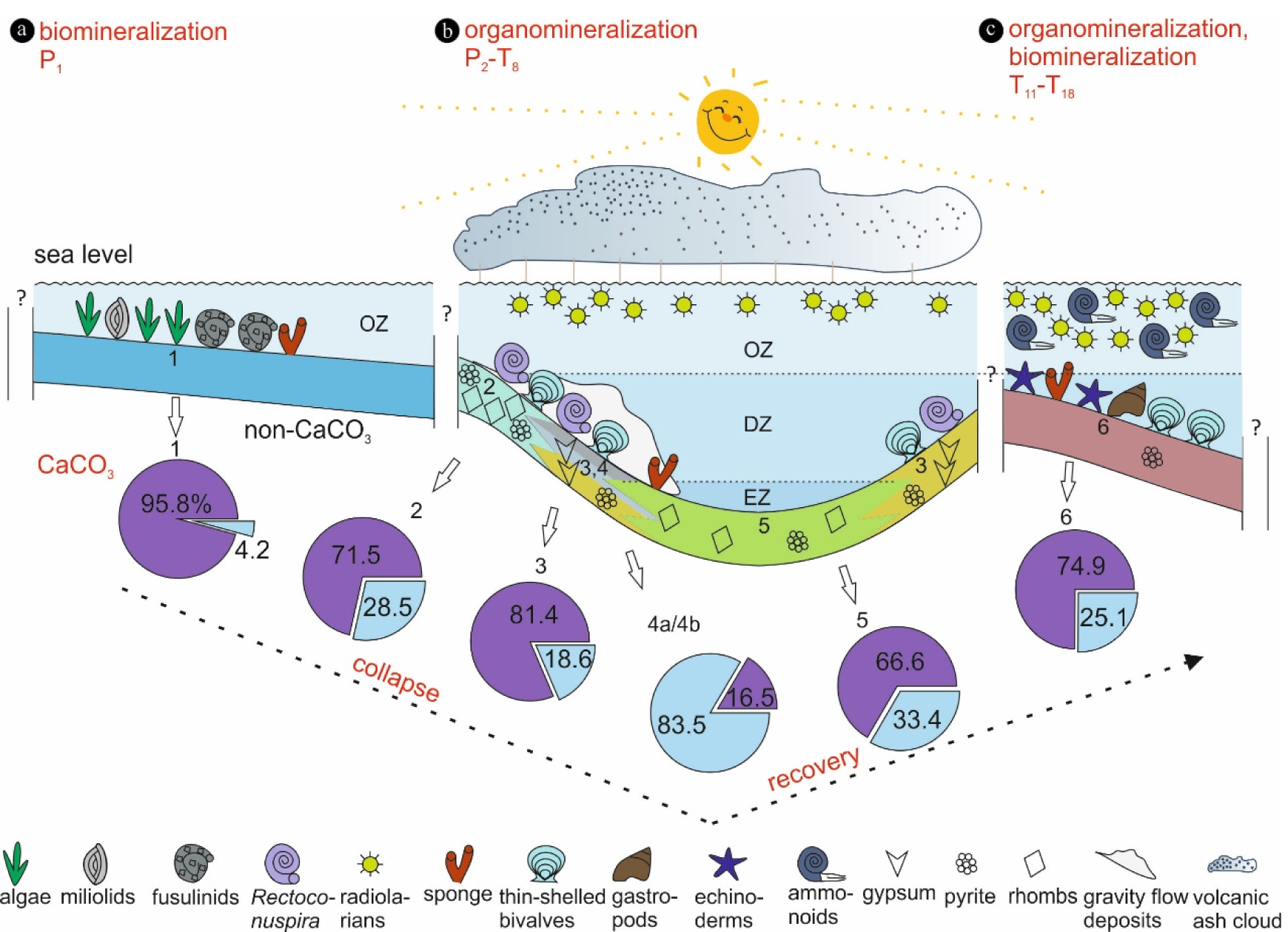
Integrative ecological model for the Xiakou area during the Permian-Triassic Critical Interval (P-TrCI). (a) Oxygenated shallow water environments and a biomineralization-dominated carbonate factory (MF-1). (b) Deeper water environments with episodically depleted oxygen, elevated salinities, and possibly euxinic conditions (MF-2–5). Carbonate was mainly formed via organomineralization (MF-2–3, 5), but this factory was intermittently interrupted (MF-4). (c) Deeper water environments and a mixed carbonate factory, with organomineralization and biomineralization being equally important (MF-6)

### 5.3 Geobiological implications for the P-TrCI

In South China, paleontological evidence and δ^13^C_carb_ data indicate the possibility of one (Jin et al. 2000; Shen et al. 2018), two (Xie et al. 2007; Yin et al. 2012; Song et al. 2013; Chen et al. 2015), or even three extinction pulses (Yang et al. 1991) in the P-TrCI. The Xiakou section records a dramatic decline in calcifying eukaryote diversity after Bed P_1_ (Figs. 2, 11). This development was accompanied by a distinct change in the carbonate factory from eukaryote-controlled to organomineralization (Fig. 12a–b). Carbonate formation by organomineralization continues until Bed P_9_, while the diversity of calcifying eukaryotes remains low. Carbonate sedimentation starts again with Bed P_16_ (Figs. 2, 12b). The diversity of calcifying eukaryotes, however, does not increase until Bed T_11_ (Figs. 2, 11, 12c). The apparent offset between carbonate sedimentation and biodiversity highlights the complexity of ecosystem change (Muscente et al. 2018) during the P-TrCI and supports the speculation that interpretations of the P-Tr extinction event might be oversimplified (cf. Erwin 1994; Stanley 2016).

At Xiakou, a dramatic decline in the diversity of calcifying eukaryotes occurred at the transition from Bed P_1_ to Bed P_2_ (Figs. 2, 11). The drop in diversity was possibly accompanied by temporary euxinic conditions, as suggested by δ^13^C_org_ signatures possibly indicative of anoxygenic phototrophs (ca. –20‰: Fig. 2; Table 1). Both developments fall into the *Clarkina changxingensis* conodont zone (see Zhao et al. 2013) and may be related to the P-Tr extinction event. At Meishan, however, the diversity of calcifying eukaryotes does not profoundly change until the uppermost part of the younger *Clarkina yini* conodont zone (Yin et al. 2001; Chen et al. 2015) (Fig. 2). Inhospitable conditions for aerobic life existed in the photic zone at Meishan for 1.5 million years prior to the widespread biological extinction, as indicated by occurrences of the lipid biomarker isorenieratane (Cao et al. 2009). The offsets in timing between the Xiakou and Meishan sections challenge the common assumption that the P-Tr mass extinction event occurred simultaneously around the globe, particularly as both sections were closely located during the P-TrCI.

## 6 Conclusions

The Permian-Triassic mass extinction is characterized by a potentially catastrophic decline of biodiversity in marine and terrestrial ecosystems. Here we presented findings from Permian-Triassic strata exposed in the Xiakou area (South China). The succession begins with a eukaryote-controlled carbonate factory (Bed P_1_). Carbonate formation by organomineralization continues until Bed P_9_. Carbonate sedimentation starts again with organomineralization in Beds P_16_–T_8_ and develops into a mixed carbonate factory, with organomineralization and biomineralization being almost equally important in Beds T_11_–T_18_. Bed P_1_ was deposited in oxygenated shallow water environments, while Beds P_2_-T_18_ were deposited in somewhat deeper environments, some of which episodically exhibited elevated salinities, oxygen depletion, and, possibly, euxinic conditions. Taken together, our studies of the sedimentary strata at Xiakou indicate an apparent disparity between carbonate sedimentation and biodiversity. Moreover, the timing of biodiversity and environmental change at Xiakou is different from that observed at the type section in Meishan, despite the proximity of the two locations. Together, these findings highlight the complexity of ecosystem change during the P-TrCI due to strong local influences, supporting the suspicion that interpretations of the P-Tr extinction event might be oversimplified.

## Acknowledgments

We appreciate constructive comments and suggestions from M. Reich, J. Peckmann and an anonymous reviewer. A. Munnecke, A.L. Claußen, D. Birgel, E. Jarochowska, G. Mathes, L.M. Baumann, M. Natalicchio, P. Suarez-Gonzalez, and S. Kiel are thanked for helpful discussions. D. Grabow, J. Luo, N. Höche, Y. He, Z-Q. Chen, H. Mei, B. Feng and G. Wu are acknowledged for field assistance. A. Hackmann, A. Pack, A. Reimer, B. Röring, C. Conradt, D. Hause-Reitner, D. Kohl, J. Dyckmans, J. Schönig, K. Lünsdorf and W. Dröse are thanked for lab assistance. This study was financially supported by the China Council Scholarship (CSC).

